# Engineering improved Cas13 effectors for targeted post-transcriptional regulation of gene expression

**DOI:** 10.1101/2021.05.26.445687

**Authors:** Emeric J Charles, Shin Eui Kim, Gavin J. Knott, Dylan Smock, Jennifer Doudna, David F. Savage

**Affiliations:** Department of Molecular and Cell Biology, University of California, Berkeley, CA, USA; Innovative Genomics Institute, University of California, Berkeley, Berkeley, CA, USA; Monash Biomedicine Discovery Institute, Department of Biochemistry & Molecular Biology; Department of Bioengineering, University of California, Berkeley, CA, USA; Molecular Biophysics and Integrated Bioimaging Division, Lawrence Berkeley National Laboratory, Berkeley, CA, USA; Gladstone Institute of Virology, Gladstone Institutes, San Francisco, CA, USA; Howard Hughes Medical Institute, University of California, Berkeley, Berkeley, CA, USA; Department of Chemistry, University of California, Berkeley, Berkeley, CA, USA; Gladstone Institute of Data Science and Biotechnology, Gladstone Institutes, San Francisco, CA, USA; California Institute for Quantitative Biosciences (QB3), University of California, Berkeley, Berkeley, CA, USA

## Abstract

Cas13 is a family of unique RNA-targeting CRISPR-Cas effectors, making it an appealing tool for probing and perturbing RNA function. However only a few Cas13 homologs have been shown to mediate robust RNA targeting in human cells, suggesting that unknown elements may be limiting their efficacy. Furthermore, many Cas13 enzymes show high degrees of toxicity upon targeting and have not been shown to mediate specific knockdown in other cell types such as *E. coli*. Here, we show that catalytically inactive Cas13 enzymes can be repurposed for efficient translational repression in bacteria with no associated growth defects. To achieve this advance, we carried out a directed evolution screen to engineer functional Cas13a variants, and identified a number of stabilizing mutations, which enabled efficient post transcriptional knockdown of gene expression. *In vitro* characterization of the resulting engineered *Lbu* Cas13a mutant, termed eLbu, revealed both stabilization and altered cleavage kinetics. Finally, we show that eLbu can be used for efficient exon skipping in human cells. This work represents the first demonstration of targeted translational repression in *E. coli* using a CRISPR enzyme, as well as the first directed evolution of a Cas13 enzyme. Such a platform could allow for engineering other aspects of this protein family to obtain more robust RNA targeting tools.

## Introduction

CRISPR-Cas systems are bacterial adaptive immune systems that protect hosts from invading foreign nucleic acids^1–4^. CRISPR loci contain integrated foreign DNA sequences (spacers) flanked by conserved palindromic repeats, which combine to express functional CRISPR RNAs (crRNAs) ^2,3^. crRNAs associate with surveillance complexes and enable a matching spacer sequence to target the effector to invading nucleic acids upon reinfection^1,2^. The rapid evolutionary arms race between bacteria and mobile genetic elements has led to the large-scale diversification of CRISPR-Cas effector functions^5^, which has been harnessed for various research and therapeutic applications^6^.

The discovery of RNA-targeting Type VI CRISPR systems has expanded the repertoire for tools to manipulate RNA^7,8^. Cas13 (formerly C2c2), is a family of sequence-activated, non-specific RNA nucleases, containing two conserved higher eukaryotic and prokaryotic nucleotide binding (HEPN) domains^7,9,10^. In contrast to other Class II systems such as Cas9, Cas13 becomes active for HEPN-mediated RNA cleavage of both cis (i.e. the target RNA) and trans (i.e. an external RNA) substrates upon sequence specific recognition of a target RNA^9,11^. The trans nuclease activity of Cas13 has initiated the field of CRISPR-based diagnostics, which has extended to DNA targeting systems with the discovery that Cas12 (formerly Cpf1) possesses a similar non-specific nuclease activity^12^. Interestingly, Cas13 has also been leveraged for targeted RNA knockdown in certain cell types, though the mechanism of cis versus trans cleavage preference remains unclear. Cas13 has also been repurposed as a targeted RNA-binding protein, via inactivation mutations to the HEPN catalytic domain^13,14^. Catalytically inactive Cas13 enzymes (ddCas13) have been shown to mediate robust transcriptome engineering (via exon skipping/inclusion), and RNA imaging in eukaryotic cells, highlighting the diversity of potential applications for an RNA-specific effector ^15^.

Due to the abundant diversity of Cas13 enzymes in nature, previous studies have relied on data mining to obtain candidate proteins and screens to validate functional variants. Such approaches, while useful in obtaining active RNA targeting enzymes, do not determine the factors necessary for an active Cas13 effector. Out of the many Cas13 sequences identified, only a handful have been shown to mediate robust RNA targeting in cells, suggesting that unknown factors may play a role in dictating their efficacy^10,16,17^. Lwa-msfGFP Cas13a, Psp Cas13b, and Rfx Cas13d (CasRx), can be repurposed for RNA knockdown in human cells, though efficiencies vary across the subfamilies^18^. Lwa-Cas13a only showed significant repression when fused to a stabilization domain, suggesting stability may be a limiting factor for this subfamily^10,13^. Furthermore, expression levels of homologs across the superfamily are variable, with Cas13a and Cas13c enzymes generally having lower levels than Cas13b for unknown reasons^19^.

Few studies have employed Cas13 enzymes for targeted knockdown in prokaryotes. In such cell types, activation of Cas13 leads to rapid non-specific cleavage of endogenous RNA, and consequently cellular dormancy^20^. Nuclease active enzymes have also been shown to lead to a high degree of cellular toxicity both in *E. coli* and native hosts, thus limiting their use as effective RNA targeting tools in such systems^10,21^. In contrast, DNA targeting tools such as Cas9 have been leveraged for targeted transcriptional knockdown via CRISPR interference (CRISPRi) in a variety of cell types^22,23^ but have associated drawbacks such as specific protospacer adjacent motif (PAM) requirements^24^.

Here we show that catalytically inactive Cas13 enzymes can be repurposed for efficient translational repression in bacteria, circumventing any toxicity associated with an active HEPN domain. Out of the initial four Cas13 homologs tested, only Cas13d led to highly efficient gene knockdown. Using directed evolution, however, we identified a series of stabilizing mutations in catalytically inactive *Leptotrichia buccalis (Lbu)* Cas13a leading to efficient translational repression in *E. coli* and RNA targeting in human cells. Stabilization of Cas13a led to altered cleavage kinetics *in vitro*, and altered binding properties, shedding light into the kinetic mechanism of Cas13a.

## Results

### Development of an effective bacterial translational repression platform using Cas13d

Analogous to transcriptional knockdown using CRISPRi in bacteria, we wondered whether catalytically inactive Cas13 enzymes (ddCas13) could be repurposed for efficient translational knockdown of bacterial mRNA when targeted to the ribosomal binding site of a target gene. We tested four catalytically inactive Cas13 homologs *Leptotrichia buccalis* (Lbu), *Lachnospiricae bacterium* (Lba), *Leptotrichia shahii* (Lsh) Cas13a, and CasRx Cas13d for knockdown in *E. coli* upon targeting the ribosomal binding sites (RBS) of a previously described endogenous dual-fluorescence RFP and GFP reporter **(Figure 1A**,**B**) ^23,25^. Notably, the synthetic RBSs of both RFP and GFP have identical sequences. Interestingly, no significant repression was observed with any dCas13a enzyme, yet dCas13d showed robust targeting, with greater than 95% fluorophore repression with both targeting guide RNAs (**Figure 1B)**.

**Figure 1.**
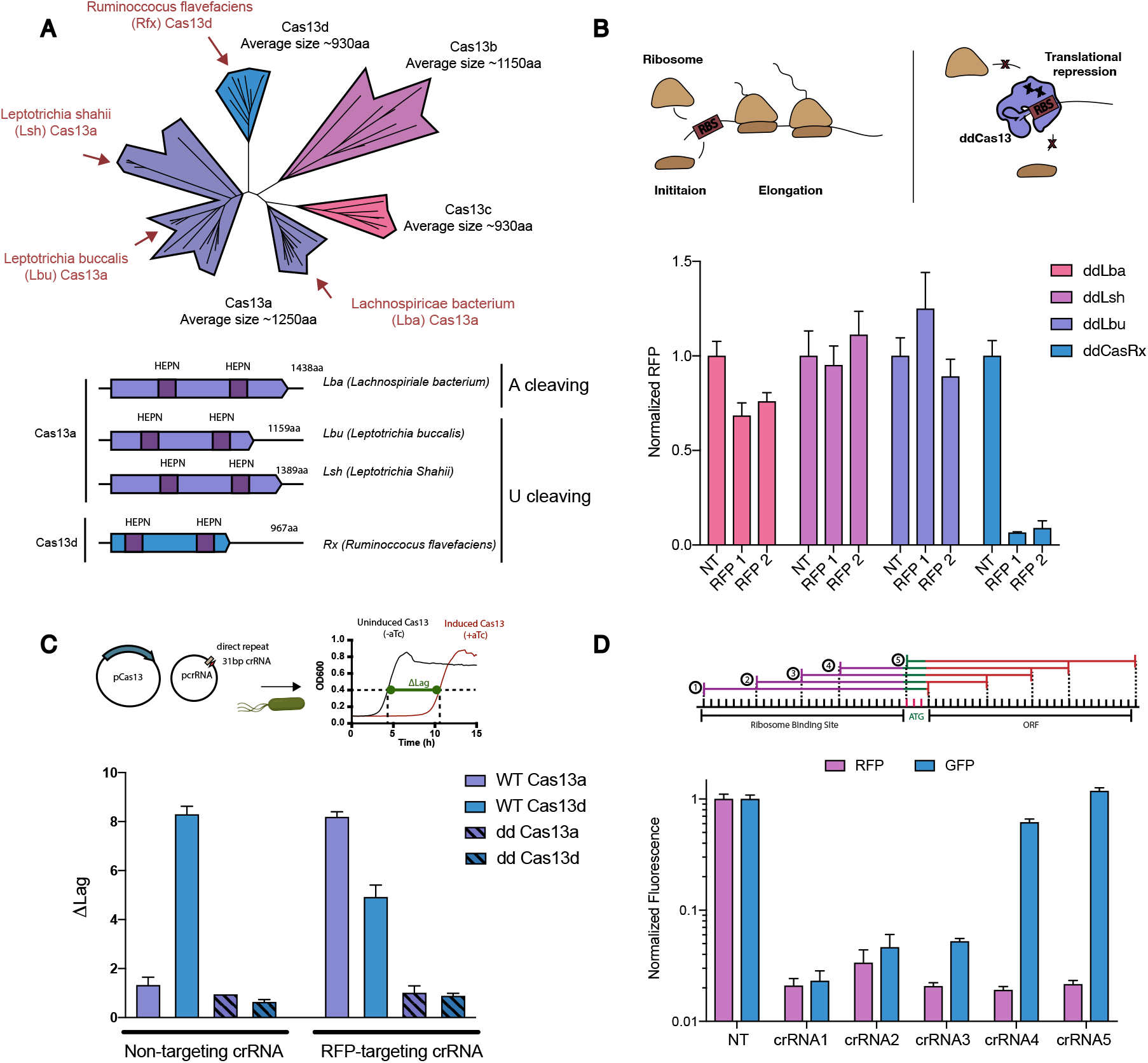
Development of an effective bacterial translational repression platform using Cas13d. **(A)** Phylogenetic tree of Cas13, with effectors used in this study highlighted in red. Below is a more detailed domain architecture of selected Cas13 enzymes. HEPN substrate preference is listed for each. **(B)** Repurposing catalytically inactive Cas13 (ddCas13) enzymes for translational repression in *E. coli*. ddCas13d efficiently represses RFP when targeted to the ribosome binding site (RBS) of RFP. Fluorescence is normalized to levels pre Cas13 induction. NT, non-targeting, RBS 1, RBS 2, two Cas13 crRNA tiling the RBS of an endogenous RFP (Values are mean +/- SEM with n=3) **(C)** Growth assays of *E. coli* transformed with either WT or catalytically inactive (dd) Cas13. Difference in lag is plotted between induced and uninduced for a non-targeting (NT) or RFP targeting (RFP) crRNA. (Values are mean +/- SEM with n=3) **(D)** RFP repression assay using ddCasRx and crRNAs tiling across the RBS of RFP, into the open reading frame (ORF). Repression can be achieved using a sequence specific to RFP only. (Values are mean +/- SEM with n=3)

To assess whether catalytically inactive Cas13 enzymes showed toxicity upon targeting in *E. coli*, a defect previously observed with Lsh Cas13, we measured the growth trajectory of cells upon induction of either WT or ddCas13. As expected, little to no growth defect was observed with any catalytically inactive Cas13 transformed with either a non-targeting or RFP targeting crRNA. In contrast, the WT version of these enzymes shows a considerable lag phase upon induction (**Figure 1C)**. Interestingly WT Cas13d showed a high degree of toxicity, even upon introduction of a non-targeting (NT) crRNA, suggesting some degree of non-specific activation of Cas13d enzymes relative to Cas13a in *E. coli*.

We next asked whether gene-specific translational repression could be obtained by targeting the beginning of the open reading frame (ORF), rather than the RBS. Five guide RNAs tiled across the RFP gene were tested for repression (**Figure 1D**). Since the RBS sequence is identical for GFP and RFP in our system, GFP fluorescence levels can be used as a control for specific targeting. All five guides showed robust RFP repression, including crRNA-5 which only targets the most 5’ end of the ORF, along with the start codon. (**Figure 1D**). GFP fluorescence was only repressed if the crRNA was complementary to the RBS, highlighting the possibility for specific translational repression upon targeting the 5’ end of an ORF.

To determine whether protein abundance could be contributing to the difference in repression between Cas13a and Cas13d, we measured protein expression by western blot against a C-terminal 2xFlag tag. Surprisingly, most Cas13a enzymes showed low levels of expression and a high amount of degradation (**Supplemental Figure 1**). In stark contrast, CasRx showed a very high degree of expression, with little proteolytic cleavage. Interestingly, Lbu Cas13 displayed a prominent truncation product at ∼100kDa. We postulated whether this was due to either proteolytic degradation or an internal RBS. Removal of the start AUG codon led to expression of a similar truncated protein (**Supplemental Figure 1**), suggesting the gene possesses an internal translational start site.

### Engineering a Cas13a based translational repression using directed evolution

Our results indicated that ddCas13a enzymes did not mediate robust RFP repression relative to the Cas13d enzyme. We therefore wondered what was limiting Cas13a in this assay and whether mutations could be identified to improve its repression activity. To this end, we generated a random mutagenesis library of Lbu Cas13a, containing >10^5^ variants with one to three mutations per gene. The library was screened for translational repression upon targeting the synthetic RBSs of both GFP and RFP (**Figure 2A**). Functional variants were isolated by fluorescence activated cell sorting (FACS) and plated on an inducer containing media. 84 single colonies exhibiting repression were picked and bulk fluorescence was measured. (**Supplemental Figure 2**). The top twelve variants were cultured overnight, miniprepped, and tested for repression upon re-transformation. All twelve variants showed robust RNA targeting, with greater than 80% fluorophore repression in bulk fluorescence experiments (**Figure 2B**). The Lbu mutagenesis library only showed significant repression when co-transfected with an RBS targeting crRNA, relative to a middle ORF targeting, or NT crRNA. (**Supplemental Figure 2**). Interestingly, western blot analysis showed that all sets of mutations increased the amount of full-length protein (**Supplemental Figure 2**), even variants containing single point mutations.

**Figure 2.**
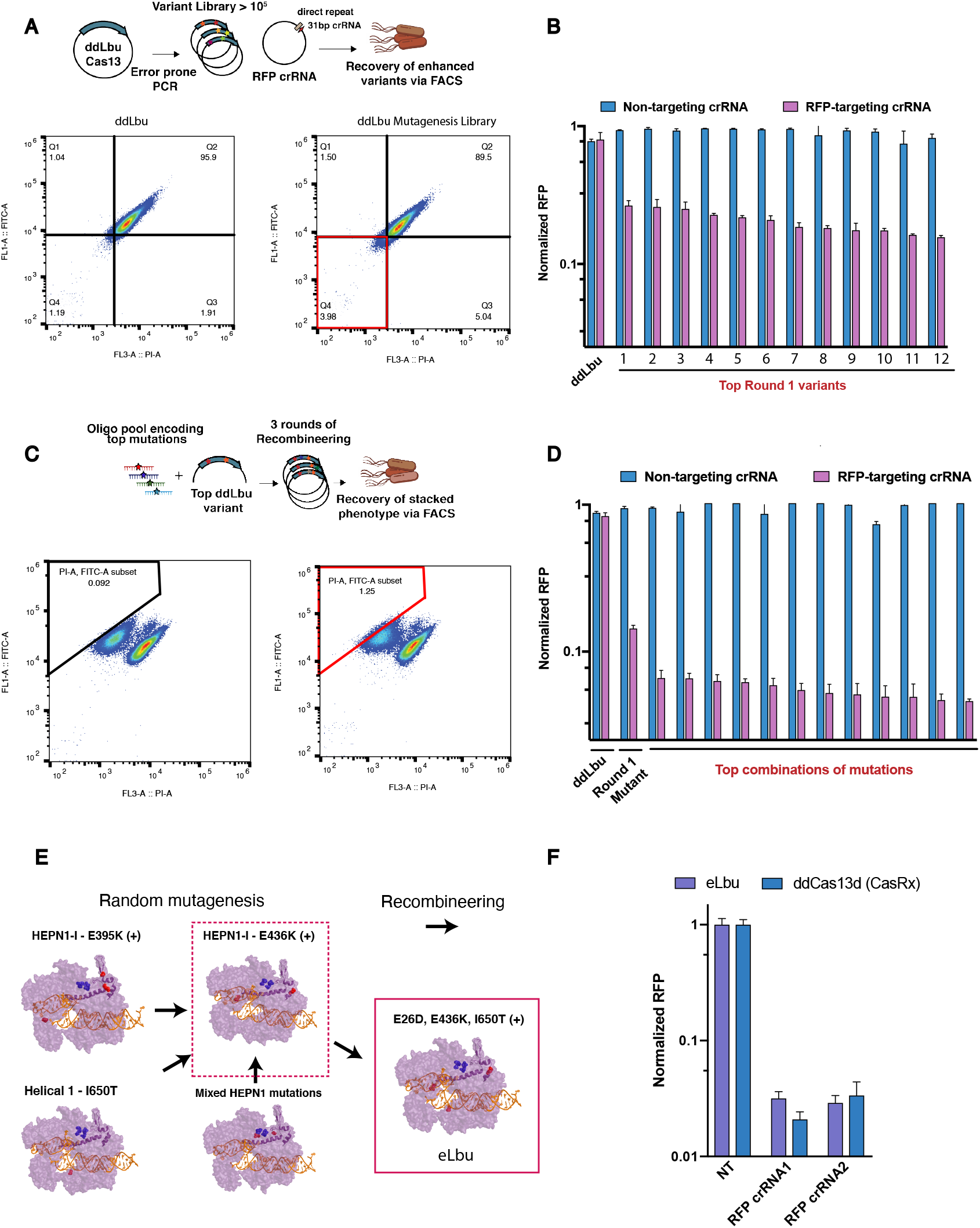
Engineering a Cas13a based translational repression using directed evolution. **(A)** Screening Lbu Cas13 random mutagenesis library for increased translational repression in *E. coli* using FACS. Only the mutagenesis library shows a decrease in fluorescence upon targeting to the RBS of RFP and GFP **(B)** Fluorophore repression of top 12 Lbu mutants following FACS sorting with a non-targeting (NT) or RFP targeting crRNA. (Values are mean +/- SEM with n=3) **(C)** Stacking top mutations from initial random mutagenesis screen using recombineering. FACS analysis shows stacking further decreases MFI of RFP. **(D)** RFP repression of top 12 Lbu stacked variants following FACS sorting. NT (non-targeting) crRNA. (Values are mean +/- SEM with n=3) **(E)** Top sets of mutations overlaid onto Lbu structure, highlighting the different classes of mutations stacked onto the E436K/E26D variants. The final eLbu variant contains three different required mutations for increased translational repression. (+) indicates that other mutations are present as well **(F)** Comparison of RFP repression using catalytically inactive eLbu and CasRx. (Values are mean +/- SEM with n=3)

Sequencing of the top 12 variants revealed that nine distinct sets of mutations in ddLbu confer efficient translational repression (**Supplemental Table 1**). Roughly one third of the variants tested contained mutations at E395, one third at E436, and one third at positions spread throughout the Helical2 and HEPN1. E395 is positioned at the end of a long helix which spans the entire HEPN1 domain, bridging the NTD and the linker domain. (**Supplemental Figure 3**). In a ternary state, E395 and E436 are positioned 3.8Å apart, but are stabilized by forming salt bridges with K433. In a binary state, these residues are greater than 6Å apart and a different stabilizing interaction is present between N440 and the peptide backbone of I393 (**Supplemental Figure 3**). Interestingly, single point mutations of E436K or E395K, alone, were not sufficient to induce efficient translational repression (**Supplemental Figure 2**), suggesting that the accompanying epistatic mutations are necessary.

To confirm that these variants were capable of programmable translational repression, the E436K E26D variant was used for knockdown of endogenous GalK, using a previously established 2-DeoxyGalactose (2DOG) negative selection system^26^. Five guide RNAs tiling the *galK* locus were tested for their ability to lead to cell survival in the presence 2-DOG (**Supplemental Figure 4**). One crRNA, in particular, led to rapid cell growth upon induction of the E435K E26D mutant, and not ddLbu (**Supplemental Figure 4**), highlighting the potential for translational repression of endogenous targets.

Next, we sought to determine whether different sets of mutations could act synergistically to increase translational repression. To this end, we used plasmid recombineering^27^ to randomly stack all 14 mutations identified from the initial screen onto the most active variant, E436K E26D (**Figure 2C, Supplemental Table 1**). The stacked variant library was targeted using an RFP specific crRNA, and variants capable of increased RFP repression were recovered via a similar FACS approach and validated by Sanger sequencing. The top engineered variants showed greater than 95% RFP repression relative to a non-targeting guide (**Figure 2D)**. Interestingly, all of the variants contained the same I650T mutation, either alone or in different genetic backgrounds, suggesting that I650T may be essential for enhanced function (**Supplementary Table 2**). E395K was not found in any variant, suggesting that E436K may be forming a salt bridge with E395 (**Supplemental Table 2**). The top performing engineered Lbu variant, termed eLbu, contained a total of seven mutations, of which E436K, E26D, and I650T are essential for efficient repression. eLbu and ddCasRx showed similar RFP repression levels of >95% when targeted to two different RBS sequences (**Figure 2F**).

### Mutations decrease Lbu’s flexibility in apo and ternary state

We wondered at what point in the enzyme kinetic mechanism these mutations might be exerting their effect. ddLbu and eLbu containing the stacked stabilizing mutations were purified and subjected to limited proteolysis when loaded with various RNA. Consistent with previously reported data for Lsh Cas13a, apo Lbu is substantially more sensitive to proteolysis than binary Lbu (**Figure 3A)**. This is most likely due to a high degree of flexibility of Cas13 in the absence of its guide RNA^28,29^. crRNA binding is known to stabilize CRISPR-Cas effector structure and likely stabilizes Lbu here, making it resistant to proteolysis treatment. Interestingly, this might be due to a non-specific electrostatic effect as addition of non-cognate RNA, such as yeast tRNA or 30 nt random RNA, also leads to a high degree of Cas13 stabilization **(Figure 3A)**.

**Figure 3.**
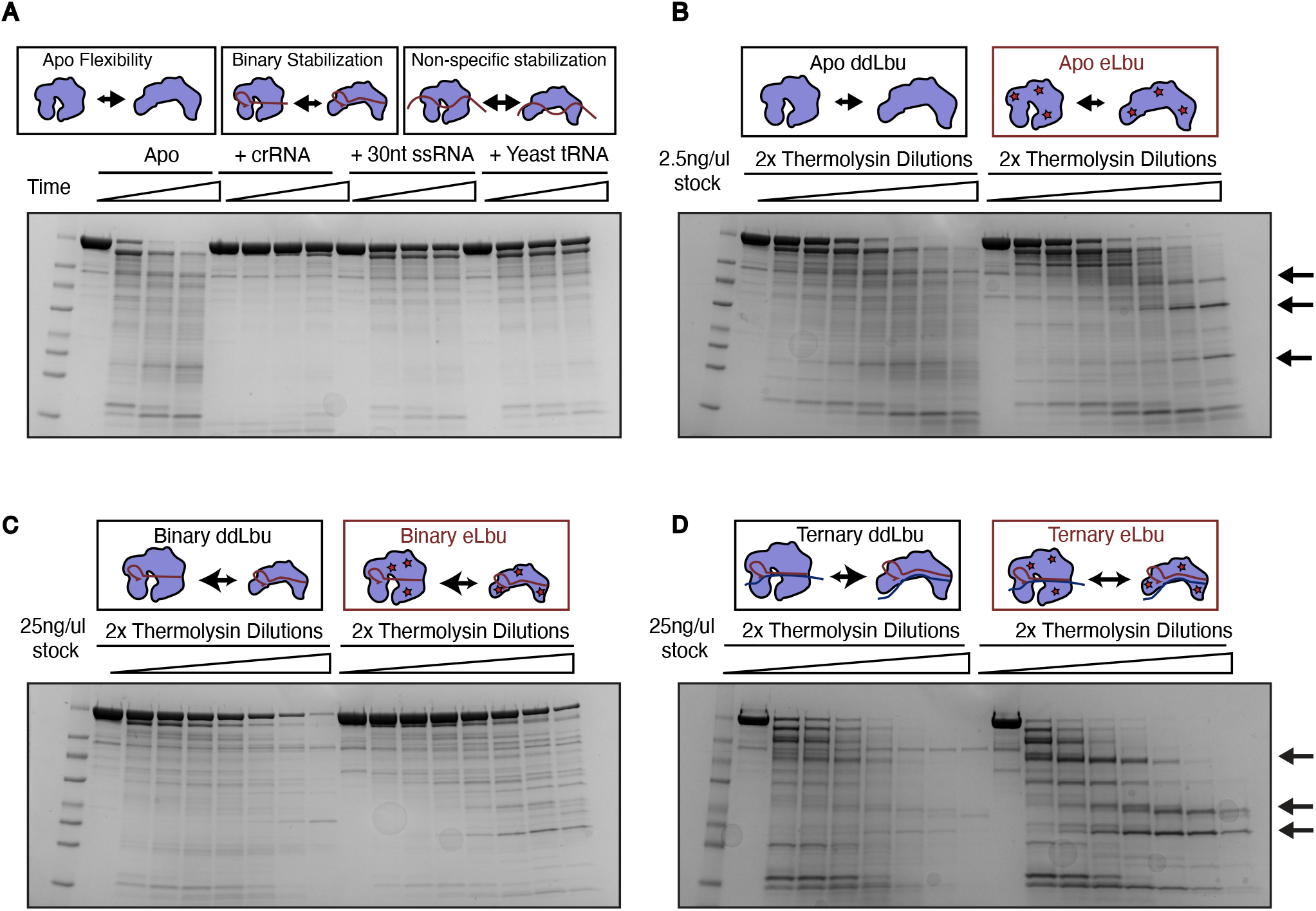
Mutations decrease Lbu flexibility in apo and ternary state. **(A)** Time points of limited proteolysis of ddLbu Cas13 using thermolysin upon addition of various RNA substrates, including cognate crRNA, 30nt ssRNA, or yeast tRNA **(B)** Limited proteolysis of ddLbu or eLbu using serial dilutions of thermolysin. Arrows indicate bands that are present in the eLbu only **(C)** Similar limited proteolysis but in the presence of 2 fold excess of crRNA. **(D)** Similar limited proteolysis experiments but in the presence of 2.5 fold excess activator RNA.

To obtain a finer resolution of proteolysis, we assayed the digestion of ddLbu and eLbu, with or without a crRNA, with increasing amounts of thermolysin. Mutations in Lbu did not increase the amount of full-length protein in the apo state but did increase the banding pattern observed at high concentrations of thermolysin (**Figure 3B**), suggesting an increase in resistance to proteolysis of the core domains. Interestingly, mutations slightly increased the amount of full-length Lbu protein in the binary state (**Figure 3C**). Similar experiments performed at 42°C also showed that mutations increase eLbu’s thermal tolerance when bound to a crRNA (**Supplemental Figure 5**).

To investigate whether mutations had any effect on the flexibility of the ternary complex, we performed the same thermolysin proteolysis assay after addition of a complementary activator RNA (**Figure 3D**). Surprisingly, activator bound Lbu showed a higher degree of proteolytic sensitivity compared to its binary state. Very little banding was observed, and the majority of the protein was degraded overtime. In contrast, eLbu showed defined banding patterns at every thermolysin concentration.

Given that mutations altered flexibility of Lbu in the apo state, we wondered whether they would also have an effect on how Cas13 interacts with non-cognate RNA. Similar thermolysin experiments were performed with addition of either a 30 nt random RNA sequence, or yeast tRNA. In contrast to ddLbu, eLbu showed lower levels of protection by either the 30 nt RNA, or the yeast tRNA, suggesting increased specificity for its cognate guide RNA (**Supplemental Figure 5**).

### Altering Cas13’s ability to associate with preformed crRNA/activator duplexes inhibits nuclease activity

We next sought to determine whether the mutations identified in ddLbu Cas13a would alter cleavage activity in the context of the wild type HEPN nuclease. To investigate this, nuclease active eLbu was purified, and tested for activity as a function of activator RNA concentration using a cleavable fluorescent reporter (**Figure 4A)**. Interestingly, reactions performed with eLbu led to early inactivation of trans cleavage at lower activator concentrations compared to WT Lbu (**Figure 4B**). While this inhibition was not observed at higher activator concentrations, eLbu’s cleavage kinetics were much slower at every concentration tested that did not saturate prior to the first measurements (**Supplemental Figure 6**).

**Figure 4.**
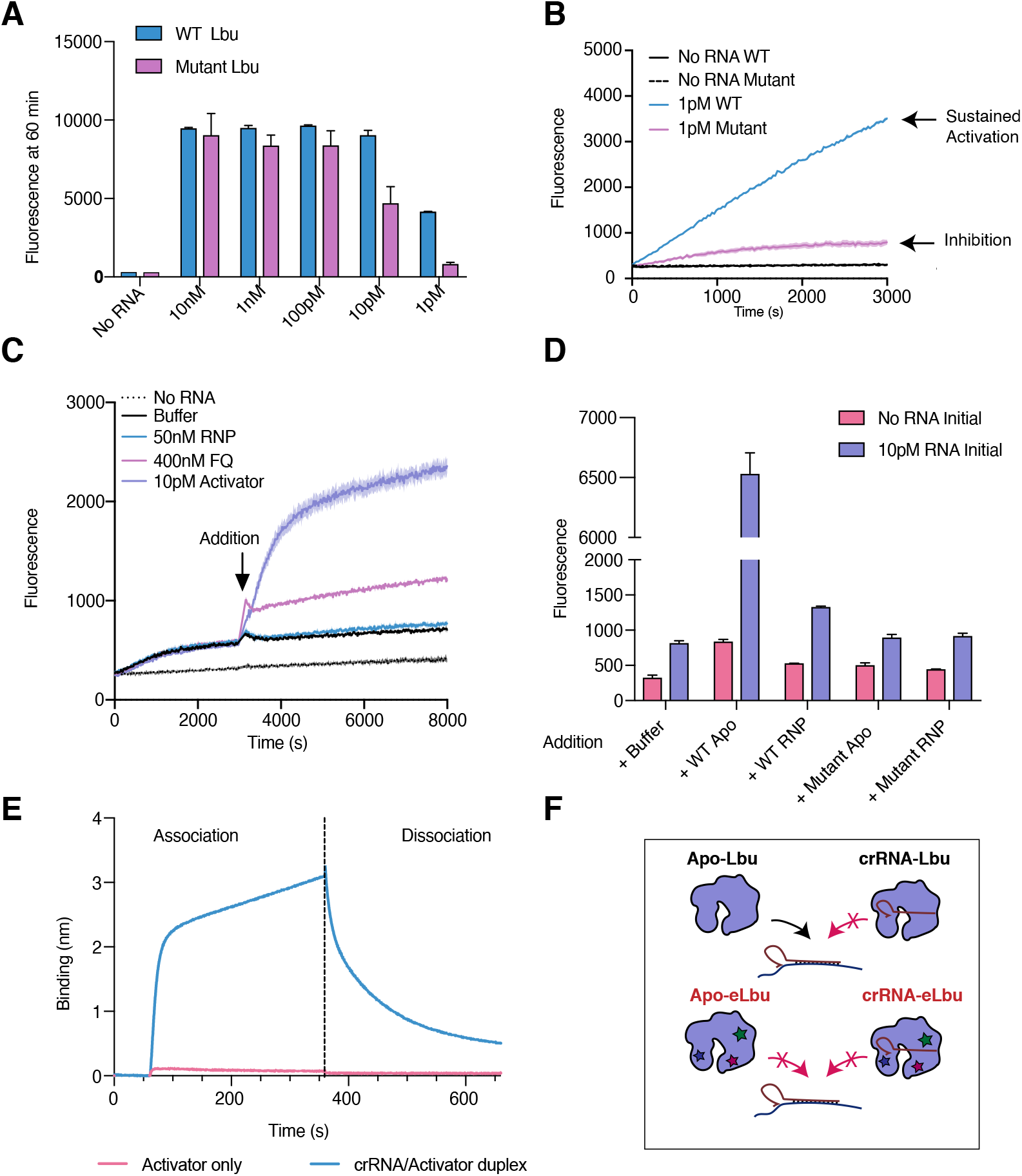
Altering Cas13’s ability to associate with preformed crRNA/Activator duplexes inhibits nuclease activity. **(A)** Cas13 activity assay using an RNA based fluorophore quencher reporter, comparing WT Lbu and mutant Lbu when titrating down activator RNA. (Values are mean +/- SEM with n=3) **(B)** Same as (A), plotted overtime at 1pM of activator RNA, comparing WT Lbu and eLbu. (Values are mean +/- SEM with n=3) **(C)** Spiking in each component into a reaction initiated with 1pM of eLbu. Only activator RNA can efficiently restart the system. (Values are mean +/- SEM with n=3) **(D)** Same as (C), but with addition of WT Lbu (apo or binary), or eLbu (apo or binary) (Values are mean +/- SEM with n=3) **(E)** BLI of apo Cas13 binding to pre-annealed crRNA/activator duplexes using biotinylated activator RNA. (Values are mean +/- SEM with n=3) **(F)** Model showing Cas13 association to preformed duplexes.

To determine what limited eLbu’s activity in the early inhibition experiments, and potentially leading to the difference in kinetics observed, each component of the reaction was re-introduced in a previously inhibited reaction. Only the introduction of activator RNA re-started the reaction, suggesting that access to activator RNA was limiting (**Figure 4C)**. This could be due to complete cleavage of the initial activator RNA by Cas13 or, alternatively, the activator could be bound to a crRNA and not available for binding. To better define the state of the activator RNA, the spike-in experiment was repeated by also testing the effect of adding either Apo or crRNA-loaded eLbu or WT Cas13. Only the introduction of Apo Cas13 was able to significantly restart the system, suggesting that the activator was present. Given the limited effect of WT RNP, it is likely the activator is already bound by an existing crRNA (**Figure 4D)**. Interestingly, eLbu in either form could not restart the reaction. This suggests that WT Cas13 may have the ability to transition directly from the apo to the ternary state, but that the eLbu mutations have altered its ability to associate with preformed crRNA-activator duplexes. To test this hypothesis, pre-annealed crRNA-activator duplexes were added to varying amounts of apo protein in the presence of a cleavable fluorescent reporter (**Supplemental Figure 7**). While incubation with WT Lbu led to high degree of fluorescence, a stark inhibition was observed with eLbu at every concentration tested, consistent with a defect in duplex association (**Supplemental Figure 7**).

To further validate that Lbu Cas13 could transition directly from apo to ternary, Biolayer interferometry (BLI) experiments were performed using immobilized biotinylated activators bound to crRNA (**Figure 4F**). As expected, WT apo Lbu showed a high degree of binding to duplexed RNA, relative to ssRNA activator alone.

### Efficient targeted exon skipping using engineered Cas13a in mammalian cells

To investigate whether the stabilizing mutations obtained in an *E. coli* system would also lead to increased function in human cells, Cas13 effectors were tested for targeted exon skipping of CD46 in HEK 293 cells (**Figure 5A**). CD46 is an endogenous, constitutively expressed cell surface receptor, present in multiple isoforms^30,31^. One isoform contains exon 12, 13 and 14, whereas another isoform only contains exons 12 and 14. Exon 13 has previously been shown to be sensitive for exon skipping via antisense oligonucleotides^29^. Using transient transfections, eLbu was targeted across exon 13 using 11 tiled crRNAs, and the amount of exon skipping was determined by RT-PCR followed by agarose gel electrophoresis (**Figure 5B**). Interestingly, many crRNAs showed a decrease in the relative abundance of long vs short isoform.

**Figure 5.**
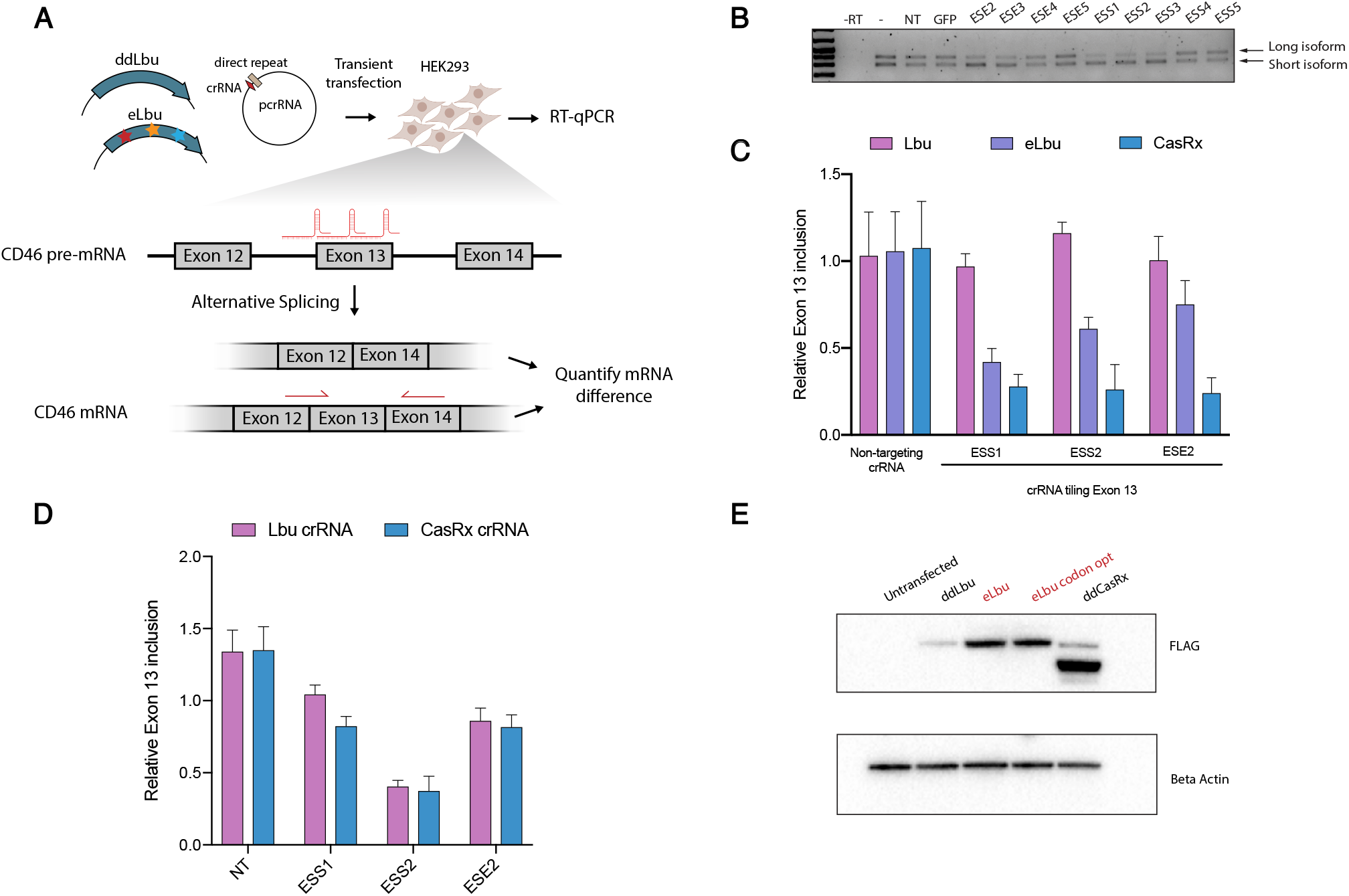
Targeted exon skipping in human cells using eLbu. **(A)** Schematic showing alternative splicing of endogenous CD46 exon13, and quantification using qPCR. **(B)** Targeting CD46 using 9 crRNAs tiling exon 13 of CD46. Relative abundance of long vs short isoform was measured using electrophoresis following RT-PCR. **(C)** Relative Exon 13 inclusion quantified by qPCR, upon targeting of ddCas13 effectors to 3 different crRNA tiling exon 13, normalized to the effects of the crRNA alone. Only CasRx and eLbu show a significant degree of exon skipping (Values are mean +/- SEM with n=3) **(D)** Relative exon 13 inclusion upon targeting of crRNA alone, comparing Cas13a and Cas13d. Transfection of crRNA alone shows a high degree of exon skipping. (Values are mean +/- SEM with n=3) (**E**) Relative expression levels of ddLbu, eLbu non codon optimized, or human codon optimized, as well as CasRx, quantified from western blot post transfection.

To further validate the amount of exon skipping, and to compare targeting across different Cas13 effectors, ddLbu, eLbu, and CasRx were targeted to CD46 pre-mRNA using the top 3 crRNA identified. The degree of isotype switching was measured by RT-qPCR using isoform specific primers. Only eLbu, as well as CasRx was able to generate >80% exon skipping with certain crRNAs, compared to ddCas13 (**Figure 5C**), when normalized to crRNA alone. Interestingly, transient transfection of crRNA alone for both Lbu Cas13a and CasRx Cas13d led to a high degree of exon skipping (**Figure 5D)**, suggesting that these short hairpin RNA may be efficiently interfering with the splicing machinery.

We then tested the expression level of ddLbu relative to eLbu upon transient transfection in HEK 293T cells, followed by western blot against a 2x FLAG tag. Surprisingly, little to no protein was present for ddLbu, when compared to the highly expressed CasRx. **(Figure 5E)**. Mutations in eLbu led to a significant 10x increase in full-length protein expression, but still remains lower than CasRx.

## Discussion

The prevalence of mobile genetic elements across the kingdoms of life has driven the evolution of a wide array of bacterial immune systems, with diverse effector functions^5^. The Cas13 family of Class II CRISPR enzymes is another unique example that has evolved to target RNA, rather than DNA. Natural Cas13 enzymes are sequence-activated, non-specific RNA nucleases, which have been leveraged for a multitude of uses, such as RNA knockdown, RNA imaging and transcriptome engineering^15–18^.

Though Cas13 enzymes have been harnessed for various human cell knock down applications, few studies have been done using Cas13 for specific repression in bacteria. This is most likely due to the high degree of toxicity associated with nuclease active enzymes. Here, we show that catalytically inactive Cas13d enzymes can be used for highly efficient translational repression upon targeting the RBS, or 5’ end of an ORF. Such a platform allows for the possibility of targeting genes at the mRNA level, and therefore within operons, something not achievable with CRISPRi. Furthermore, the lack of PAM requirement of Cas13 enzymes dramatically increases the potential target sites within a given sequence, with all crRNAs tested showing a high degree of repression.

While CasRx Cas13d natively possesses the capability for efficient translational repression, all Cas13a enzymes tested showed little to no knockdown. To this end, we performed the first high throughput mutagenesis screen on Lbu Cas13a, combining both random mutagenesis and targeted plasmid recombineering, to identify a multitude of variants capable of efficient translational repression in *E. coli*. The top variant also showed robust exon skipping capabilities in human cells, suggesting that mutations were transferable across various cell types. The observed amenability of Cas13 for mutagenesis could pave the way for engineering other aspects of this protein, such as increased on-target cleavage. Furthermore, two orthogonal translational repression platforms could be useful for developing CRISPR based synthetic gene circuits.

Cas13 has been shown to induce cellular dormancy of its native host upon phage infection, rather than cell death^20^. The reversible nature of this dormancy implies some type of inactivation mechanism during Cas13’s kinetic mechanism, which has yet to be identified. Here, we show that Cas13 can transition directly from the apo state to the ternary state, if given a preformed crRNA-activator duplex. Altering the flexibility of Cas13 in the apo state via mutations not only altered its ability to associate to pre-formed duplexes, but led to a stark difference in cleavage kinetics, as well as an early inhibition of cleavage activity at lower concentrations *in vitro*. This suggests that under normal conditions, Cas13 may be cycling on and off a preformed crRNA-duplex once the initial single-stranded activator binding occurs. In such a scenario, inactivation could occur if Cas13 is given a competing RNA to bind to, such as free crRNA. This could provide a natural off switch in Cas13 once enough phage RNA becomes cleared, and the ratio of free crRNA to duplexed crRNA/target RNA increases.

Proteolytic regulation has been shown to control various stress response pathways in bacteria^32^. Akin to the changes in sigma factor stability observed during stress, the various degrees of flexibility and proteolytic susceptibility demonstrated by Cas13 could have been driven by evolutionary pressure to shut off activated Cas13 at a certain point in its kinetic cycle. The high degree of proteolytic cleavage observed in the ternary state could either be due to the flexible nature of activator bound Cas13, or due to release of crRNA-activator duplexes by Cas13, and return to its apo state. In either case, once a complementary target RNA is present, Cas13 will be activated for non-specific RNA cleavage, but will rapidly become degraded, thus depleting the pool of active Cas13, while leaving the binary RNP surveillance population intact. More studies looking at the effects of Cas13 mutations on their ability to lead to dormancy could shed light on the contribution of proteolytic turnover to Cas13 inactivation.

Overall, our data indicates that catalytically inactive Cas13 enzymes can be used for robust translational repression in *E. coli*. Using directed evolution, we engineered two orthogonal platforms for gene knockdown, which could be used for high throughput screens or metabolic engineering in prokaryotes. Mutations obtained in an *E. coli* system were transferable to mammalian cells, highlighting the strength of such a platform for engineering other aspects of this protein. Altering Cas13’s ability to associate to preformed crRNA/duplexes also led to slower cleavage kinetics, suggesting the potential for Cas13 to be cycling on and off an activator RNA.

## Methods

### Plasmid cloning

For *E. coli* constructs, Lbu, Lsh, and Lba Cas13a sequences were PCR amplified from previously described vectors^11^ and cloned into Tetracyclin inducible backbones containing chloramphenicol resistance markers. CasRx sequence was amplified from addgene plasmid pXR001 and cloned in identical vectors. crRNA for Cas13a, PCR amplified from previously described vectors^11^, and CasRx amplified from addgene pXR003, were cloned into ampicillin containing backbones, inserted between the crRNA handle and terminator. For mammalian cell expression, the same sequences were cloned into transient expression vectors containing EF1a promoter, and puromycin resistance gene. crRNA were expressed from the same plasmid under U6 promoter and polyT terminator using previously described vectors^33^.

### *E. coli* translational repression

Plasmids encoding catalytically inactive Cas13 under inducible Tet promoter, and constitutively transcribed crRNA were cotransfected in *E. coli* MG1655 harboring genomically encoded RFP and GFP as previously described^23,25^. Briefly, single colonies were picked and cultured to log phase in a 96 deep well plate, using Teknova EZ media at 37°C. Cultures were then diluted 1:1000 into +/- 20nM aTc containing media, and fluorescence was measured after 6h of growth in a Tecan M1000.

### Random mutagenesis library and FACS analysis

Lbu Cas13 was amplified using Agilent GeneMorph II random mutagenesis kit with varying ratios of input DNA to cycle number to obtain a mutation rate of 1-3 per gene. PCR amplicon was ligated into Tet inducible backbone via golden gate and transformed in Top10 cells to obtain >10^5^ colonies. Mutagenesis library was grown for 8h, miniprepped, and cotransfected with a NT or targeting crRNA into MG1655 harboring genomically encoded RFP and GFP^23^, and cultured to log phase. Cultures were then diluted 1:1000 into 20nM aTc containing media, and incubated at 37°C for 6h, and fluorescence was analyzed via FACS. Fluorescence was gated using the non-targeting library as a negative control. Cells were then plated on +/- 20nMaTc containing LB-agar plates, and incubated at 37°C overnight. 100 colonies exhibiting low levels of fluorescence were picked and cultured overnight in the absence of aTc. Cultures were then assayed for translational repression as described above.

### Plasmid recombineering and FACS analysis

Plasmid recombineering was performed as described previously^27^. Briefly, oligonucleotides encoding the top twelves mutations were electroporated into induced ECNR2 strain alongside a plasmid encoding the top Cas13 variant identified from the mutagenesis screen. Cultures were grown overnight at 37°C in LB media, mini prepped, and subjected to two more rounds of recombineering in a similar fashion. Two separate reactions were carried through all three rounds, one using no input DNA, and one using the 12-oligonucleotide library. Screening recombineering library for translational repression was done as described above but using an RFP specific crRNA.

### Protein purification

Protein purification was performed as described previously^11^. Briefly, cultures containing MBP-His-Tev-Cas13 were induced with 0.5mM IPTG at mid log phase, following overnight incubation at 16°C. Cells were then pelleted, and lysed using sonication. Clarified lysate was loaded on a Ni-Nta column, washed, and eluted in high Imidazole. Cas13 protein was incubated with TEV protease, dialyzed overnight, and further purified using a combination of ion exchange and gel filtration.

### Limited proteolysis assays

Purified Cas13 was incubated at a final concentration of 1uM with varying concentrations of thermolysin (2µg/µl-0.03µg/µl) in reaction buffer (150mM KCl, 20mM Tris, 1mM TCEP, 10mM MgCl_2_) for 30 min at 37ºC. When indicated, Cas13 was incubated with 2x molar ratio of crRNA, and 2.5x molar ratio of activator RNA. Reactions were then inactivated at 95ºC for 5 min, run on SDS-PAGE and stained by Coomassie.

### *In vitro* fluorescent cleavage assay

Cas13 was first complexed with crRNA at 1:1 ratio for 10 minutes at room temperature in reaction buffer (50mM KCl, 20mM HEPES pH 6.8, 5mM MgCl_2_, 5% Glycerol) at a final concentration of 500nM. Reactions were then diluted to a final concentration of 50nM in the presence of varying concentrations of activator RNA, and 400nM polyU fluorescent reporter. Fluorescence was measured in a Tecan Spark at 37ºC over the course of 2 hours. For spike in experiments, reaction was left to saturate in the Tecan for 30min, after which 2ul of each component was spiked in.

### Transient transfection of human cell lines

HEK293 cells were cultured in DMEM Glutamax containing 10% FBS. 20k cells were seeded in a 96 well plate 24h before transfection. Plasmids were transfected in triplicate with lipofectamine 3000 using 100ng of DNA. Cells were selected 24h post transfection with puromycin at a final concentration of 1µg/ml for 48h.

### RNA extraction and qPCR

RNA was harvested by phenol-chloroform extraction, combined with ethanol precipitation, and reverse transcribed using Ecodry RNA to cDNA premix containing random hexamers. cDNA was then amplified using Dynamo HS SYBR green with CD46 isoform specific primers, and normalized to primers targeting an upstream region of CD46. Ct counts were normalized to either untransfected cells, or cells transfected with crRNA alone.

## Acknowledgements

This work was supported by NIH grants 1R01GM127463 (D.F.S) and RM1HG009490 (J.A.D, D.F.S), by Defense Advanced Research Projects Agency (DARPA) award HR0011-17-2-0043 (J.A.D) and by Mammoth Biosciences (D.F.S). We thank Ben Oakes, Sean Higgins, Rob Nichols, Luke Oltrogge, and Avi Flamholz for technical support and productive discussions.

## Declaration of Interestsxs

J.A.D. is a co-founder of Caribou Biosciences, Editas Medicine, Intellia Therapeutics, Scribe Therapeutics, and Mammoth Biosciences. J.A.D. is a scientific advisory board member of Caribous Biosciences, Intellia Therapeutics, eFFECTOR Therapeutics, Scribe Therapeutics, Synthego, Metagenomi, Mammoth Biosciences, and Inari. J.A.D is a member of the board of directors at Driver and Johnson & Johnson. D.F.S. is a co-founder of Scribe Therapeutics and a scientific advisory board member of Scribe Therapeutics and Mammoth Biosciences. All other authors declare no competing interests.

