## Supplemental Figures and Tables for "Engineering improved Cas13 effectors for targeted post-transcriptional regulation of gene expression"

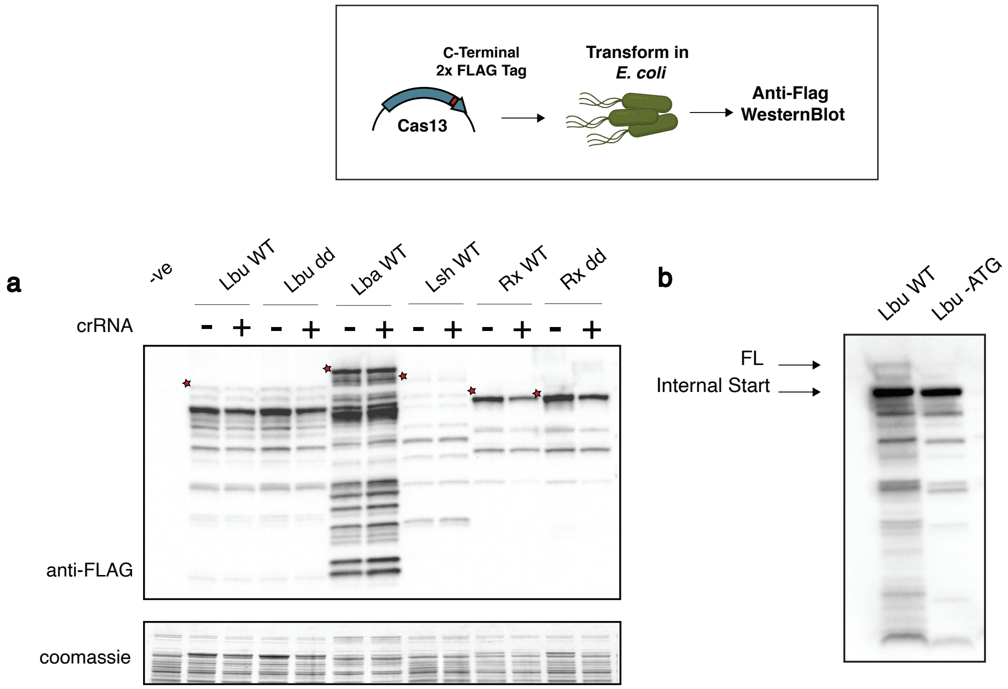


**Supplemental Figure 1 – Screening Cas13 expression in *E. coli* across Cas13a and Cas13d orthologs**

(A) Western blot analysis comparing various Flag-tagged Cas13 effectors, in the presence or absence of a cognate crRNA. Constructs were transformed in *E. coli,* and induced for 3h at 37C, following anti-FLAG western blotting. (B) Similar western blot as in (A), looking at the effect of removing the start codon from WT Lbu. The persistence of the smaller band shows that this product is due to an internal RBS, leading to a truncated product.

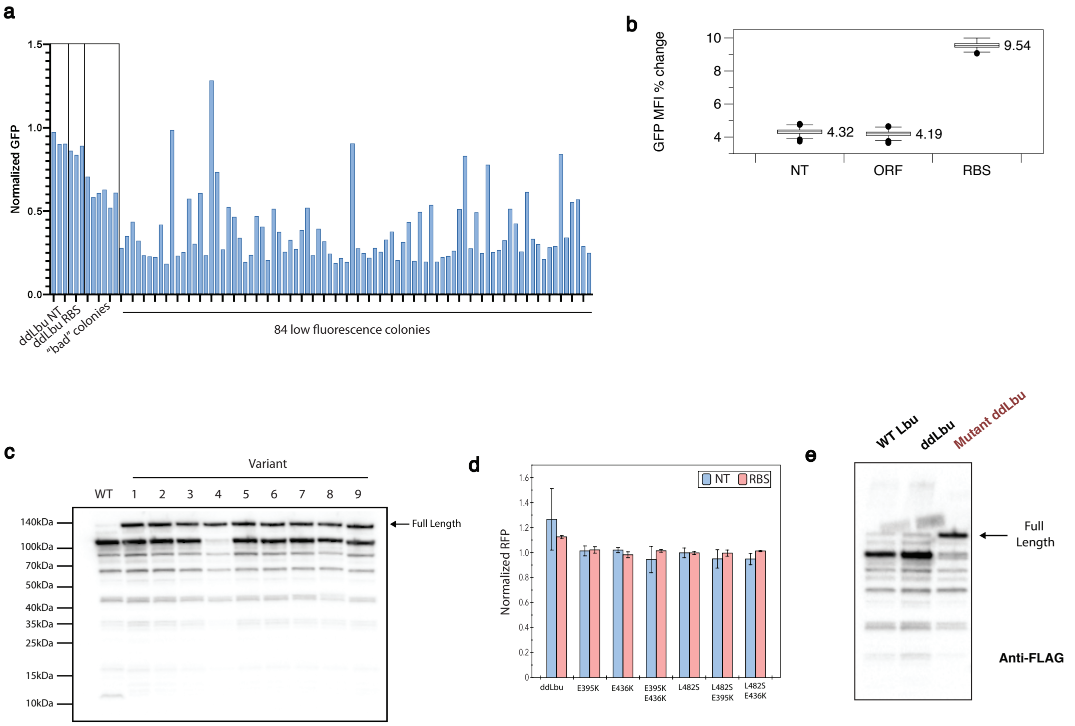


**Supplemental Figure 2 – Screening ddLbu random mutagenesis library for increased translational repression**

(A) GFP repression of individual ddLbu random mutagenesis variants post FACS sort. Colonies were plated on 20nM aTc containing media plates. Functional repressing colonies were picked by visualizing the plate under blue light. The top twelve variants were mini prepped and retransformed for downstream validation. (B) Bootstrap analysis of the mean fluorescence intensity change between ddLbu and the ddLbu random mutagenesis library co-transformed with either a non-targeting (NT), open reading frame (ORF) or ribosome binding site (RBS). Only targeting to the RBS shows a change in the MFI. (C) Anti-flag western blot analysis showing increased expression of each variant from the random mutagenesis screen. (D) RFP repression assay showing that single point mutations E395K and E436K alone do not confer efficient translational repression. (E) Anti-flag western blot of the top performing variant, eLbu, showing increased expression compared to either WT or ddLbu

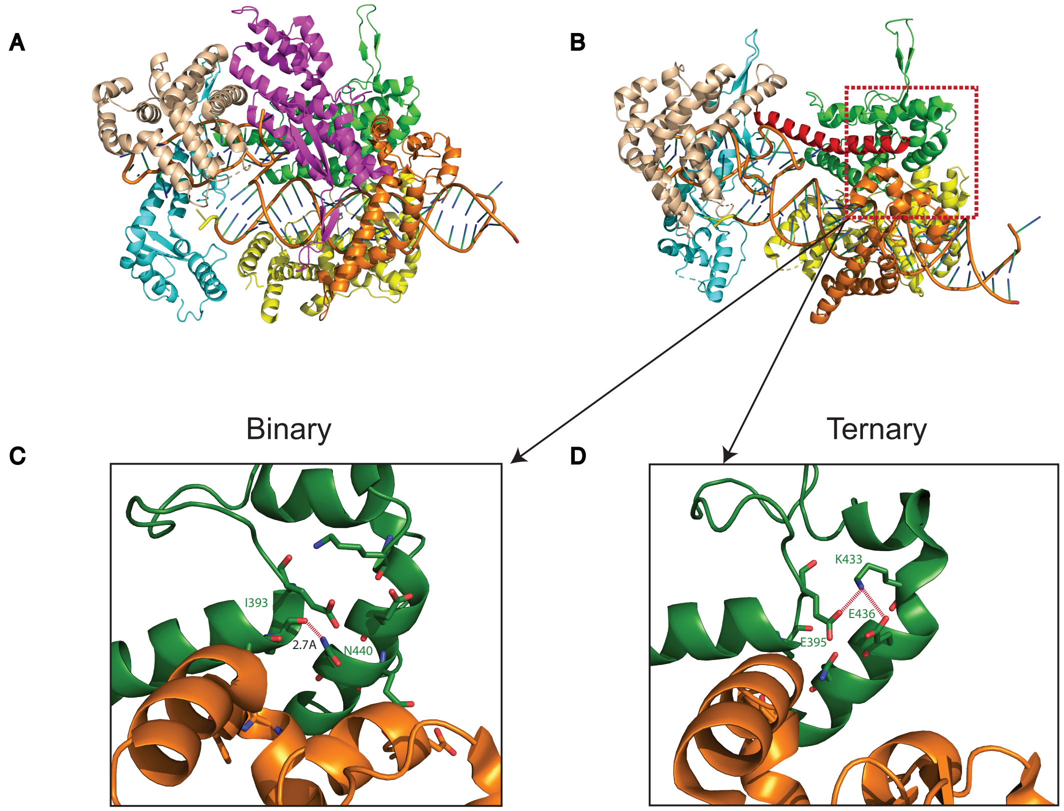


**Supplemental Figure 3 – E395 and E436 are coordinated in a ternary state**

(A) Ternary ddLbu Cas13 complex (PDB:5XWY) with each subdomain colored. (B) Same as (A) with the HEPN2 domain removed. The long helix spanning HEPN1 is colored in red. (C) Zoom into the hinge region where E395 is located in a binary state. N440 is salt bridged with I393 and E395/E436 are far apart. (D) Same as (C) but in a ternary state. E395, E436 and K433 are involved in a network of salt bridge interactions

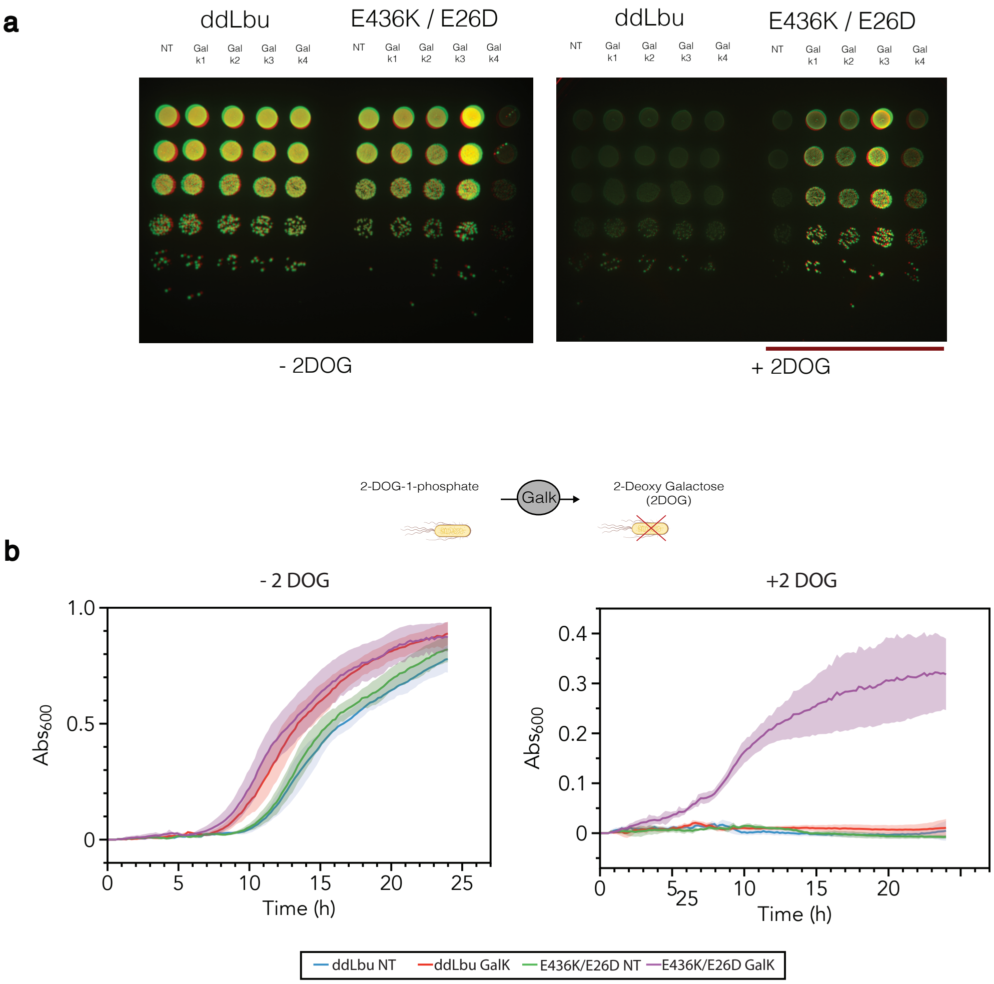


**Supplemental Figure 4 – Efficient endogenous GalK translational repression using engineered Lbu Cas13**

(A) Spot plating assay screening four GalK targeting crRNA for viability upon addition of 2-DOG. RFP/GFP containing *E. coli* were used for this assay. Addition of 2DOG to ddLbu cotransfected with guides led to small off-white colonies. In contrast, targeting GalK using E436K/E26D Lbu led to large colonies which maintained both RFP and GFP fluorescence. (B) Growth assay using the top GalK targeting crRNA comparing ddLbu and E436K/E26D ddLbu. Only E436K/E26D ddLbu showed growth if co-transfected with a GalK targeting crRNA.

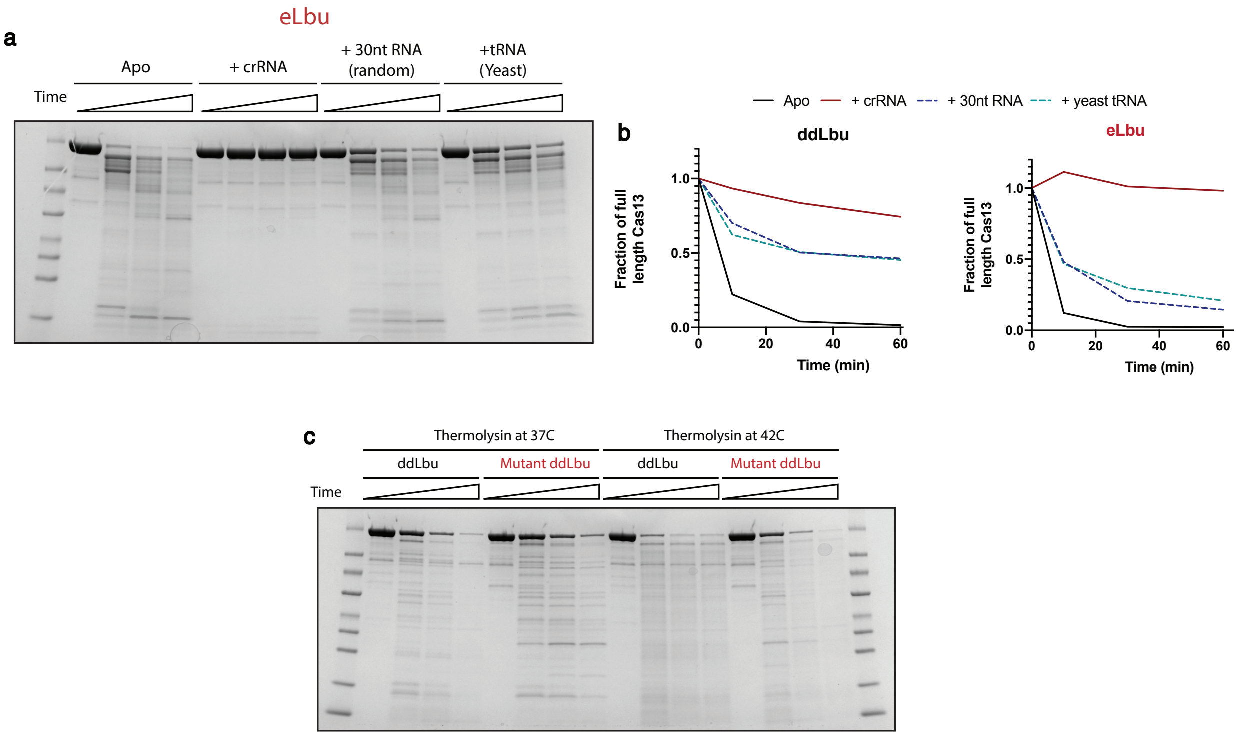


**Supplemental Figure 5 – eLbu is more specific for cognate crRNA, and more thermostable**

(A) Limited proteolysis using thermolysin at different time points (5 min, 15min, 30min) using purified eLbu Cas13 complexed with cognate crRNA, 30nt ssRNA, or yeast tRNA. (B) Quantification of full length Cas13 from the same experiments comparing ddLbu with catalytically inactive eLbu. (C) Thermolysin experiments using crRNA bound Cas13 at 37C, or 42C, comparing ddLbu with eLbu at different time points. A small increase in full length protein is observed with eLbu at 42C, compared to ddLbu.

Supplemental Figure 6

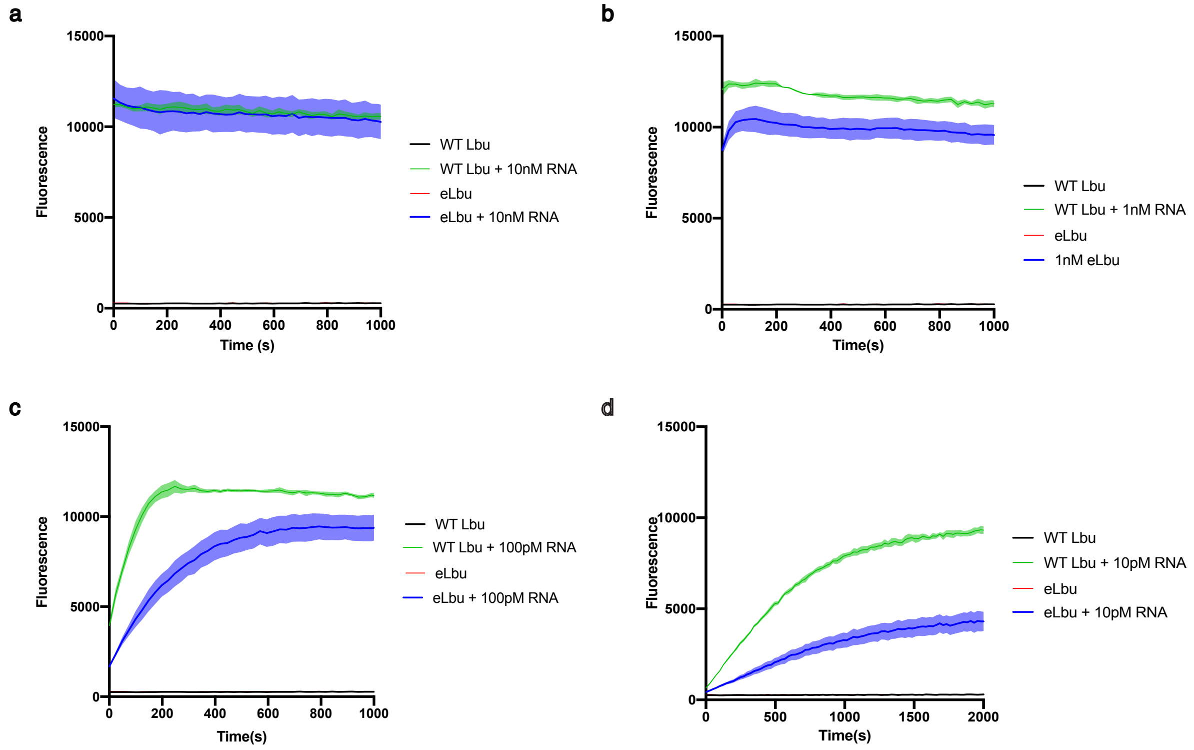


**Supplemental Figure 6 – eLbu Cas13 shows slower cleavage kinetics in fluorophore/quencher assay**

RNAse detection assay comparing 50nM WT Lbu to nuclease active eLbu at (A) 10nM activator RNA, (B) 1nM activator RNA, (C) 100pM activator RNA, (D) 10pM activator RNA. eLbu in both (C) and (D) showed much slower cleavage kinetics compared to WT Lbu.

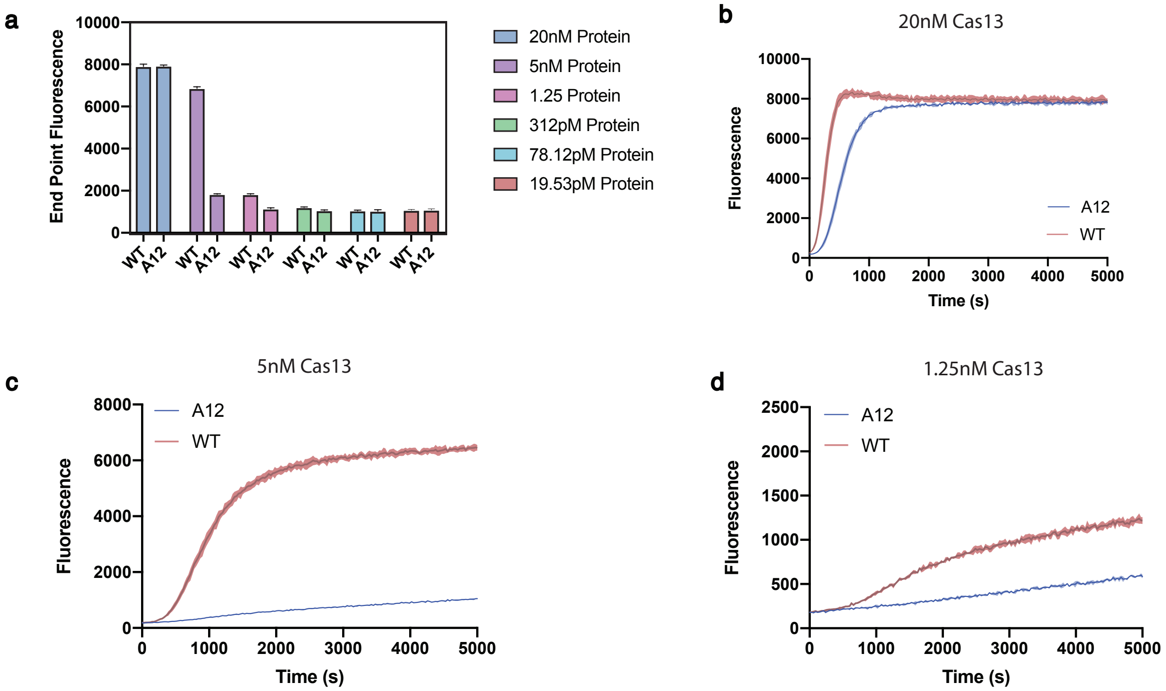


**Supplemental Figure 7 – Cas13 activation using preannealed crRNA activator duplexes**

(A) Cas13 endpoint activity assay comparing WT Lbu to nuclease active eLbu at different protein apo concentrations, using 5nM activator RNA preannealed to 10nM crRNA. A significant hit in activity is observed for eLbu below 20nM Cas13. (B) Same as (A) but measuring activity overtime at 20nM Cas13. (C) 5nM Cas13 (D) 1.25nM Cas13

| Variant# | Frequency | Mutations |
|---|---|---|
| 1 | 3 | L218S, S422Y, E395K |
| 2 | 2 | E26D, E436K |
| 3 | 1 | E436K, E709K |
| 4 | 1 | D91E, M264I, E436K, N568D |
| 5 | 1 | E395G, A472D |
| 6 | 1 | L482S, A494V |
| 7 | 1 | L753P |
| 8 | 1 | N796E |
| 9 | 1 | I650T |
| **Total** | **12** | |

**Supplemental Table 1 – Top 9 variants identified from the ddLbu random mutagenesis screen**

Mutations were identified by sanger sequencing the top twelve variants selected following FACS sorting and plating of the ddLbu random mutagenesis library. E395 and E436 appear mutated in multiple instances, as well as in different genetic backgrounds.

| Variant# | Frequency | Mutations |
|---|---|---|
| 1 | 1 | D91E, S422Y, L482S, A494V, N568D, I650T |
| 2 | 1 | I650T,Q796E |
| 3 | 1 | <u>L218S,M264I,L482S,I650T,L753P</u> |
| 4 | 1 | I650T,L753P,A472D |
| 5 | 1 | M264I, L482S, A494V, N568D, I650T |
| 6 | 1 | L218S, M264I,I650T,L753P |
| 7 | 1 | M264I, I650T |
| 8 | 1 | M264I, I650T,L753P |
| 9 | 1 | L218S,M264I,L482S,I650T |
| 10 | 1 | D91E,L218S,M264I,N568D,I650T |
| 11 | 1 | D91E, S422Y, I650T |
| 12 | 1 | I650T |

**Supplemental Table 2 – Top 12 variants identified from the recombineering screen**

Mutations were identified by sanger sequencing the top twelve variants selected following FACS sorting and plating of the ddLbu recombineering library. Every variant contains I650T in different genetic backgrounds, as well as one variant that contains I650T alone. The eLbu variant selected for downstream applications is underlined.
